# Derivation, Characterizations, and Applications of Rabbit Haploid Embryonic Stem Cells

**DOI:** 10.1101/2025.11.05.686447

**Authors:** Zhenpeng Zhuang, Manya Yu, Tao Cui, Tong Qiu, Yuanxi Yang, Longquan Quan, Lei Li, Quanjun Zhang, Yinghua Ye, Zhen Ouyang, Chengcheng Tang, Junwei Wang, Junjie Mao, Liangxue Lai

**Affiliations:** Institute of Laboratory Animal Sciences, Chinese Academy of Medical Sciences & Peking Union Medical College, Beijing, 100730, China; Guangdong Provincial Key Laboratory of Stem Cell and Regenerative Medicine, Guangzhou Institutes of Biomedicine and Health, Chinese Academy of Sciences; China-New Zealand Joint Laboratory on Biomedicine and Health, Guangzhou Institutes of Biomedicine and Health, Chinese Academy of Sciences; Sanya institute of Swine resource, Hainan Provincial Research Center of Laboratory Animals, Sanya 572000, China; Guangdong Provincial Key Laboratory of Large Animal models for Biomedicine, Wuyi University, Jiangmen 529020, China; University of Chinese Academy of Sciences, Beijing 100039, China; School of Biology and Brewing Engineering, Taishan University, Tai’an 271018, PR China; Analytical Instrumentation Core, Guangzhou Institutes of Biomedicine and Health, Chinese Academy of Sciences

**Author notes:** These authors contributed equally (Zhenpeng Zhuang, Manya Yu).

## Abstract

Haploid embryonic stem cells (haESCs), which contain a single set of chromosomes, provide powerful systems for investigating gene function and creating gene-edited animal models. haESCs have been derived from several mammalian species including human, mouse and rats, yet not from rabbits. Here, we report the derivation of parthenogenetic haESCs of rabbit (rbPhESCs) using an optimized culture medium. These cells maintain a stable haploid karyotype during long-term culture and own pluripotency features comparable to diploid embryonic stem cells derived from rabbit fertilized embryos. By integrating gene trapping cassette into *ROSA26* locus, rbPhESCs offer an ideal platform for high-throughput functional genomic screening, as shown in haESCs from other mammals. Consistent with their parthenogenetic origin, rbPhESCs durably preserve maternal-specific imprinting patterns even after extended culture in serum-containing conditions, providing a valuable platform for generating disease models deficient in maternally expressed imprinted genes. Therefore, our findings expand the repertoire of mammalian haESCs and establish rbPhESCs as a valuable platform for genetic studies and biotechnological applications in rabbits.

## Introduction

In most organisms, diploidy is the default state of somatic cells, while haploidy occurs only in specific biological contexts. Haploid cells, which carry a single set of chromosomes, naturally arise in certain species such as honeybees and ants [1], and in vertebrates they are confined to the post-meiotic germ line, including sperm and oocytes [2]. Fortunately, studies have demonstrated that haploid cells can be transiently maintained in androgenetic, gynogenetic, or parthenogenetic blastocysts [3,4]. Building on this observation, and with the rapid progress in embryonic stem cell (ESC) technologies, the derivation of haploid embryonic stem cells (haESCs) in vertebrates has become feasible. To date, haESCs have been successfully established in several species, including medaka fish, mice, rats, sheep, cows, monkeys, and humans, largely owing to the development of optimized culture conditions [5–11]. However, the derivation of haESCs in many other species remains a considerable challenge.

Compared with diploid embryonic stem cells (diESCs), haESCs share many characteristics, including the capacity for indefinite self-renewal and the ability to differentiate into derivatives of all three germ layers [12]. Specifically, haESCs can form embryoid bodies in vitro and give rise to teratomas when injected subcutaneously into immunodeficient mice, both of which contain numerous terminally differentiated cell types [13]. Moreover, the germline competence of haploid ESCs has been demonstrated by tracking coat color and transgenic markers through the female germline, indicating that cells cultured with a haploid genome retain germline transmission potential similar to that of diESCs [14].

However, haESCs also possess unique features that enable applications not attainable, or more effective, than those achievable with conventional diploid systems [15]. Notable applications include generation of bi-maternal or bi-paternal offspring [16], construction of stable karyotypes in individuals through programmed chromosome fusion [17], production of allodiploid cells [18], gene trapping based high throughput screening [19], complete telomere-to-telomere genome sequence assembling [20,21], and elucidation of interactions between inter- and intra-chromosomal elements of individual mammalian genomes [22], establishing haESCs as a valuable cellular resource.

Furthermore, the single set of chromosomes in haESCs is inherited exclusively from either the maternal or paternal genome, classifying them as parthenogenetic haESCs (phESCs) or androgenic haESCs (ahESCs) [23], both of which can function as gametes and give rise to fertile offspring [24–26]. Remarkably, bimaternal and bipaternal offspring were successfully generated using haESCs with deletions in specific imprinted regions, providing a powerful strategy for generating animal models with targeted, parent-of-origin–specific genetic modifications [16,27].

Rabbits are important not only as farm animals in agricultural production but also as valuable models in life science research, contributing significantly to our daily life and the development of therapeutic drugs [28]. In recent years, several key milestones have further strengthened the role of rabbits in biological research, including the establishment of germline-competent pluripotent cell lines [29], the generation of single-cell transcriptomic datasets [30], and comparative analyses across species [31–34]. These advancements have facilitated a wide range of studies in rabbits, including, but not limited to, antibody structure [35], vaccine production [36,37], the modeling of human diseases [38], and developmental biology [39,40]. Unlike rodents, whose embryos develop as egg cylinders, rabbit embryos form bilaminar discs during gastrulation, a developmental mode shared with humans and most other mammals [41]. This similarity, together with practical advantages such as a relatively short gestation period, the production of multiple offspring per pregnancy, and delayed implantation following the onset of gastrulation [42], makes rabbits particularly valuable for the direct visualization of early developmental events.

Given the versatile applications of haESCs and the growing importance of rabbits in biomedical research, establishing haESCs in rabbits will be of great significance. Here, using an optimized culture medium, we report for the first time the successful derivation of rabbit parthenogenetic haploid embryonic stem cells (rbPhESCs) from parthenogenetically activated embryos. Although only a small fraction of haploid cells was present in the newly derived rbPhESCs, they could be stably maintained at a high proportion after successive rounds of enrichment by fluorescence-activated cell sorting (FACS). Cytogenetic analysis confirmed that rbPhESCs possess 22 chromosomes, representing half of the normal chromosome number in rabbits. Similar to diESCs of rabbit (rbDiESCs), these cells exhibit hallmark features of pluripotency, including high alkaline phosphatase activity, embryoid body formation, and teratoma development. Transcriptome profiling by RNA sequencing (RNA-Seq) further revealed that rbPhESCs express core pluripotency genes and exhibit a transcriptional profile corresponding to an earlier developmental stage than that of E6.5 primordial germ cells (PGCs). Importantly, we demonstrate that rbPhESCs are amenable to high-throughput genetic screening and can serve as effective donors for semi-cloning, highlighting their potential as a valuable platform for functional genomics in rabbits and the generation of imprinting-related disease models.

## Results

### Establishment of Rabbit Haploid Parthenogenetic Embryonic Stem Cells (rbPhESCs) from Rabbit Blastocysts

In mammals, haploid embryonic stem cells could be derived from blastocysts resulted from either in vitro activated mature oocytes or zygotes with removal of the paternal or maternal pronucleus. Based on this principle, we sought to establish rbPhESCs by using blastocysts achieved from parthenogenetically activated oocytes (Fig. 1A). Since a mucinous layer often exists around rabbit embryos, particularly when flushed from the oviduct at later stages (Fig. S1A), in order to facilitate ICM hatching (Fig. 1A), we introduced a small nick in the zona pellucida (ZP) before plating blastocysts onto feeder cells (Fig. S1A). Expectedly, Compact, colony-like outgrowths emerged from ZP-nicked blastocysts approximately six days after plating and were subsequently isolated for further propagation (Fig. S1B).

**Figure 1.**
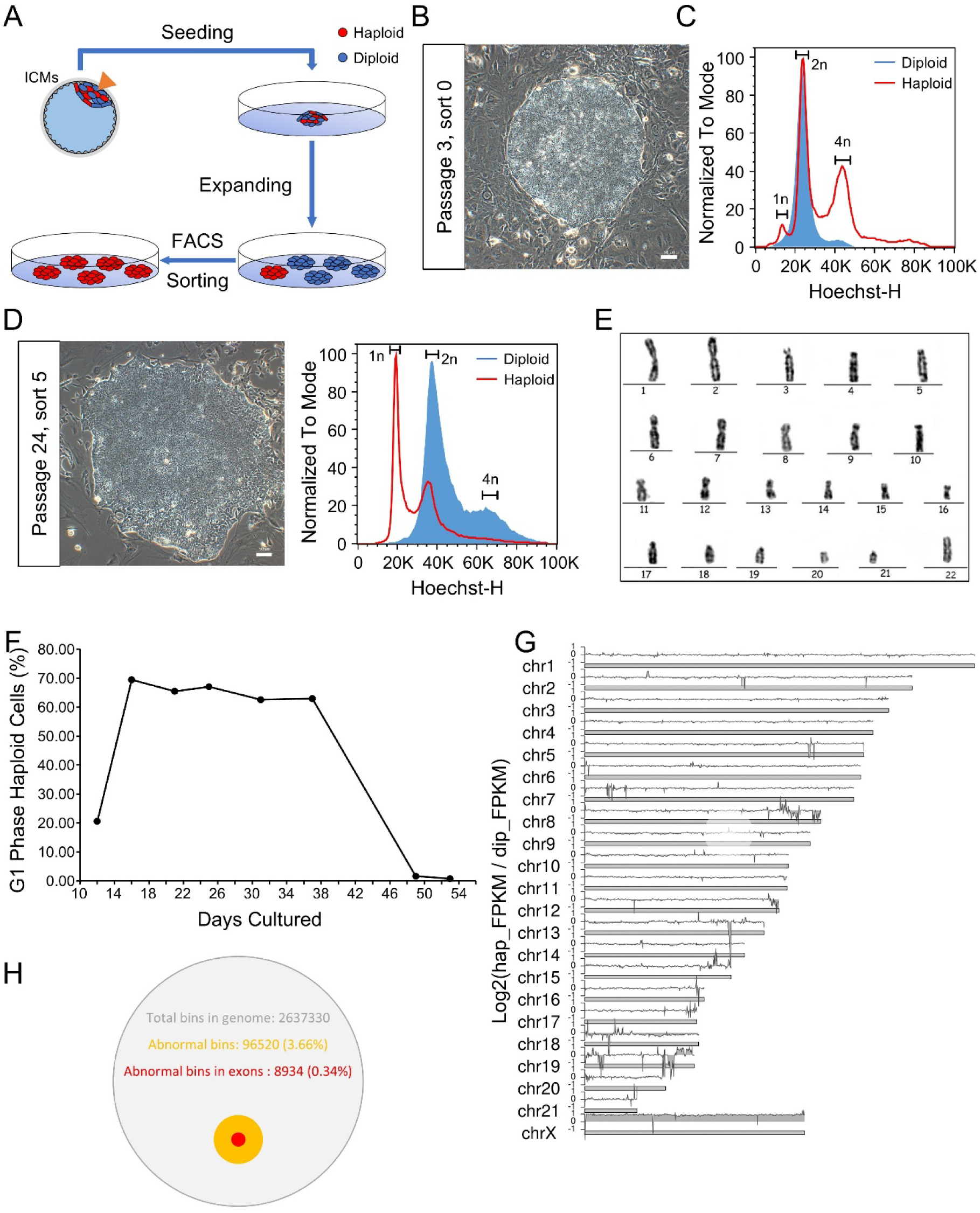
Derivation and stable culture of rabbit parthenogenetic embryonic stem cells (rbPhESCs) **(A)** Schematic diagram of the derivation of rabbit haploid embryonic stem cells. The zona pellucida near the inner cell masses (ICMs) of E4.0 parthenogenetic blastocysts was mechanically nicked to facilitate cell outgrowth, and the embryos were immediately plated onto feeder cells. After several passages for expansion, the cells were dissociated and stained with 5 µg/mL Hoechst for fluorescence-activated cell sorting (FACS) to isolate haploid cells. The sorted haploid cells were then replated onto feeders for further proliferation. (**B**) Colony morphology of rbPhESCs prior to sorting, proliferating on feeder cells. Scale bar: 100 µm. (**C**) Distribution of cells with different DNA contents during the first round of sorting. Cells were stained with 5 µg/mL Hoechst 33342 for DNA content analysis.1n: haploid cells in the G0/G1 phase; 2n: haploid cells in S phase or diploid cells in G0/G1 phase; 4n: diploid cells in S phase. (**D**) Phase-contrast image of rbPhESCs after 24 passages of continuous culture on feeder cells, following completion of five rounds of haploid cell sorting. Scale bar: 50 µm. DNA content of rbPhESCs at the sixth round of FACS analysis. 1n, 2n, and 4n peaks represent the same cell cycle phases as described in panel C. (**E**) Karyotype analysis of rbPhESCs at passage 25, numbers under the chromosomes are randomly assigned, not represent the number of chromosomes. (**F**) Percentage of haploid rbPhESCs in G1 phase during continuous in vitro culture, cells in G1 phase were identified and quantified by FACS analysis. (**G**) Copy number variation analysis of rbPhESCs. Rectangles represent chromosomes, and the broken lines above them indicate the log2-scaled copy number ratio comparing rbPhESCs to diploid cells. Hap_FPKM: FPKM-normalized read counts within 100,000 bp bins for rbPhESCs; Dip_FPKM: FPKM-normalized read counts within 100,000 bp bins for male diploid rabbit ESCs. (**H**) Proportion of genomic bins in rbPhESCs showing copy number changes compared to diploid ESCs. The green circle indicates the percentage of rbPhESC bins with more than a twofold change in copy number relative to diploid ESCs. The yellow circle represents the percentage of exonic regions within these altered bins.

After several passages, rbPhESCs derived from the initial outgrowth displayed a compact colony morphology in condition of both with and without feeders (Fig. 1B, S1C, S1D). A major challenge in haESC derivation is spontaneous diploidization that occurs during prolonged culture. In our study, we observed that, initially, only a small fraction of cells in rbPhESCs were haploid (<0.5%) (Fig. 1C). During passaging to expand the cell population, we performed fluorescence-activated cell sorting (FACS) every two weeks to enrich haploid populations. After five rounds of enrichment, almost 100% cells were haploid and exhibited typical colony morphology (Fig. 1D).

Karyotype analysis revealed that rbPhESCs possessed a haploid chromosomal set (Fig. 1E, Fig. S1E), which was stably maintained for over 38 days of continuous culture without FACS-based enrichment (Fig. 1F). Moreover, de novo whole-genome sequencing revealed that rbPhESCs retained complete genomic sequences comparable to those of male diploid rbESCs (Fig. 1G). Only 3.66% of genomic regions displayed copy number variations greater than twofold, with just 0.34% located within exonic regions (Fig. 1H).

In summary, rbPhESCs have been successfully established from rabbit parthenogenetic activated blastocysts and their compact colony morphology and haploid chromosome set are able to remain stable even after long term in vitro culture.

### rbPhESCs Exhibited Pluripotency Similar to Rabbit Diploid Embryonic Stem Cells (rbDiESCs)

A recent study demonstrated that rbDiESCs, either derived from fertilized blastocysts or reprogrammed from somatic cells, can be maintained in a primed pluripotent state [29]. Consistently, our rbPhESCs exhibited similar characteristics to those two types of diploid cells, including the expression of core pluripotency markers *OCT4*, *SOX2*, and *NANOG* (Fig. 2A), along with strong alkaline phosphatase (AP) activity (Fig. 2B). When cultured in suspension without KOSR, CHIR99021 and IWR-1, rbPhESCs formed typical embryoid bodies (EBs) (Fig. 2C, S2A). Furthermore, subcutaneous injection of rbPhESCs into severe combined immunodeficiency (SCID) mice, teratomas with a size of around 2 cm formed in eight weeks and contained differentiated derivatives of all three germ layers (Fig. S2B, 2D).

**Figure 2.**
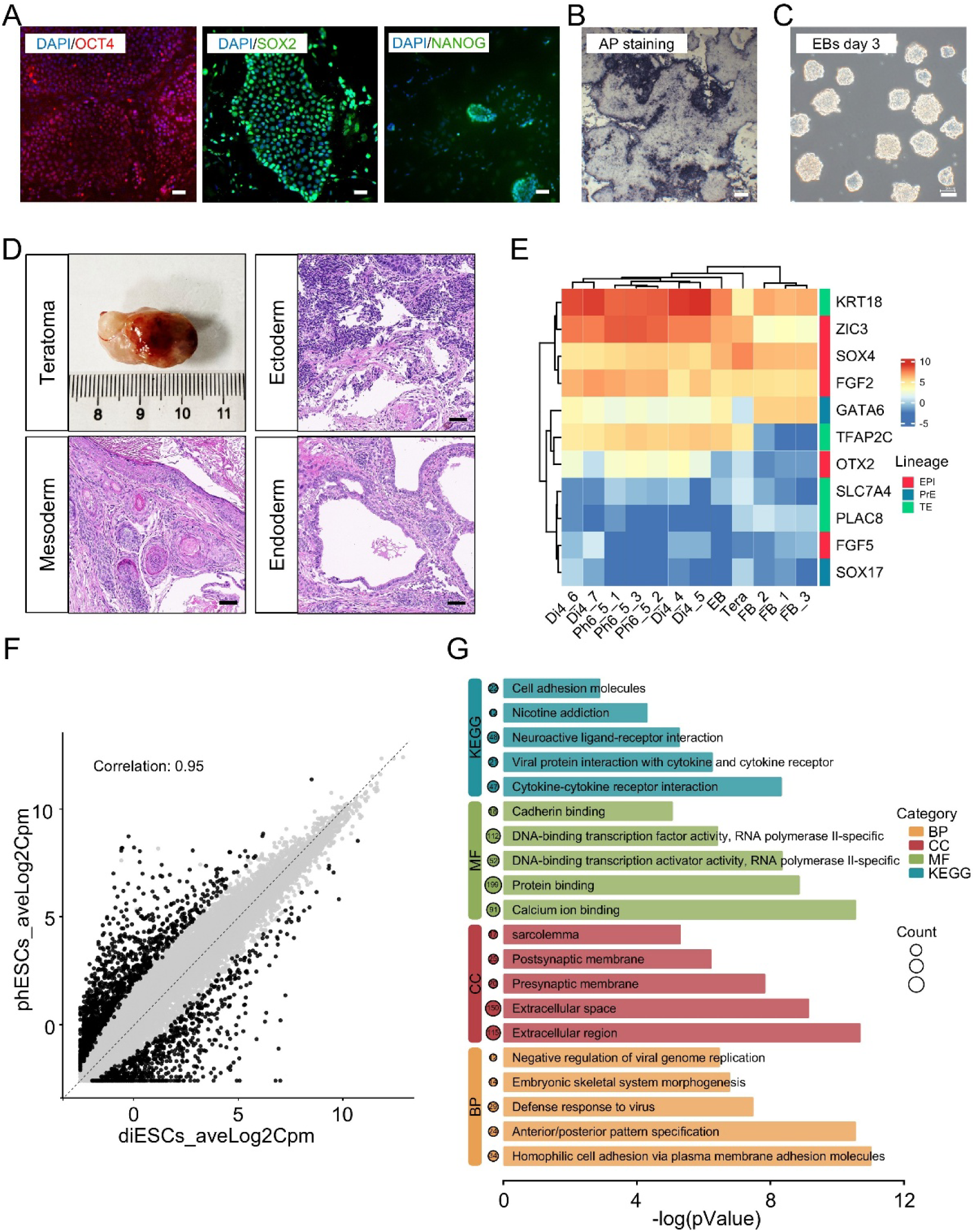
Pluripotency and Transcriptome Analysis of rbPhESCs. **(A)** Immunofluorescence images showing expression of pluripotency markers in rbPhESCs. Nuclei are counterstained with DAPI (blue). Scale bar: 50 µm. (**B**) Alkaline phosphatase (AP) staining of exponentially proliferating rbPhESCs cultured on feeder cells in iKFC medium. Scale bar: 10 µm. (**C**) Phase-contrast images of embryoid bodies (EBs) formed after 4 days of differentiation. rbPhESCs were dissociated into single-cell suspensions and cultured under suspension conditions in iKFC medium lacking CHIR99021 and IWR-1. Scale bar: 50 µm. (**D**) Image of teratocarcinoma (top-left) and H&E staining sections illustrating representative derivatives of the three germ layers. Scale bar for the sections: 50 µm. (**E**) Hierarchical clustering of samples based on expression of lineage-specific genes for epiblast (EPI), primitive endoderm (PrE), and trophectoderm (TE). Di: rabbit diploid embryonic stem cells; Ph: rabbit parthenogenetic embryonic stem cells; EB: embryonic body; Tera: teratocarcinomas; FB: fibroblasts (**F**) Global transcriptomic correlation analysis between rbPhESCs and rabbit diploid embryonic stem cells (rbDiESCs). Raw gene expression counts were normalized using log-transformed counts per million (CPM). Each sample included three biological replicates, and the final CPM value for each gene represents the average across the three replicates. (**G**) Gene ontology (BP: Biological Process; CC: Cellular Component; MF: Molecular Function) and pathway enrichment (KEGG) analysis of significantly differentially expressed genes between rbPhESCs and rbDiESCs.

In addition, we compared the transcriptomic profiles between rbPhESCs and rbDiESCs generated under identical culture conditions though RNA sequencing. Rabbit somatic cells (fibroblasts from one-month-old rabbits), and differentiated derivatives (rbPhESCs-derived EBs and teratomas) were used as control. Hierarchical clustering of global gene expression patterns revealed that rbPhESCs clustered closely with rbDiESCs, while distinctly from somatic and differentiated samples (Fig. S2C). To further define their developmental identity, we examined lineage-specific markers associated with the epiblast (EPI), primitive endoderm (PrE), and trophectoderm (TE) [43]. Our data showed that rbPhESCs and rbDiESCs exhibited highly similar expression patterns across all three lineages. This resemblance was reinforced by direct transcriptomic comparison, which showed a strong correlation between the two populations (Fig. 2F). Notably, as rbDiESCs, rbPhESCs also expressed *OTX2*, a marker of the formative/primed pluripotent state [44] (Fig. 2E). It was true that a small number of genes were differentially expressed between the two types of the stem cells, while they were enriched in pathways and Gene Ontology terms unrelated to core pluripotency networks (Fig. 2G). Collectively, these results demonstrate that rbPhESCs and rbDiESCs share a highly similar molecular identity and confirm that rbPhESCs possess pluripotency equivalent to that of rbDiESCs.

### rbPhESCs maintained the maternal genomic imprinting patterns

In addition to exhibiting pluripotency comparable to diploid ESCs, oocyte-derived haploid embryonic stem cells have been shown to preserve parental-specific genomic imprinting at the *H19*–*IGF2* and *GTL2*–*DIO3* loci [45]. For *H19* gene, due to lack of detailed annotation in rabbits, we first identified the putative H19 transcripts using our RNA-Seq data. As showed in Fig. S3A, the uncharacterized ncRNA LOC127486901 (Gene ID: 127486901), located between *IGF2* and *RPL23*, exhibited high expression in rbPhESCs, but was nearly undetectable in rbDiESCs, indicating a maternal-specific expression pattern. (Fig. 3A). We therefore considered this ncRNA as the *H19* of rabbits. We then analyzed its evolutionary relationship among other mammals. Phylogenetic analysis based on sequences from 17 mammalian species revealed that the rabbit *H19* gene shares a closer evolutionary relationship with the human counterpart than with those of other mammals, except nonhuman primates (Fig. 3B). For *IGF2*, a paternally expressed imprinted gene, we compared its functional domains with mouse. As showed in Fig. S3B, strong conservation across nearly all functional domains was observed, while its expression was nearly absent in rbPhESCs (Fig. 3A).

**Figure 3.**
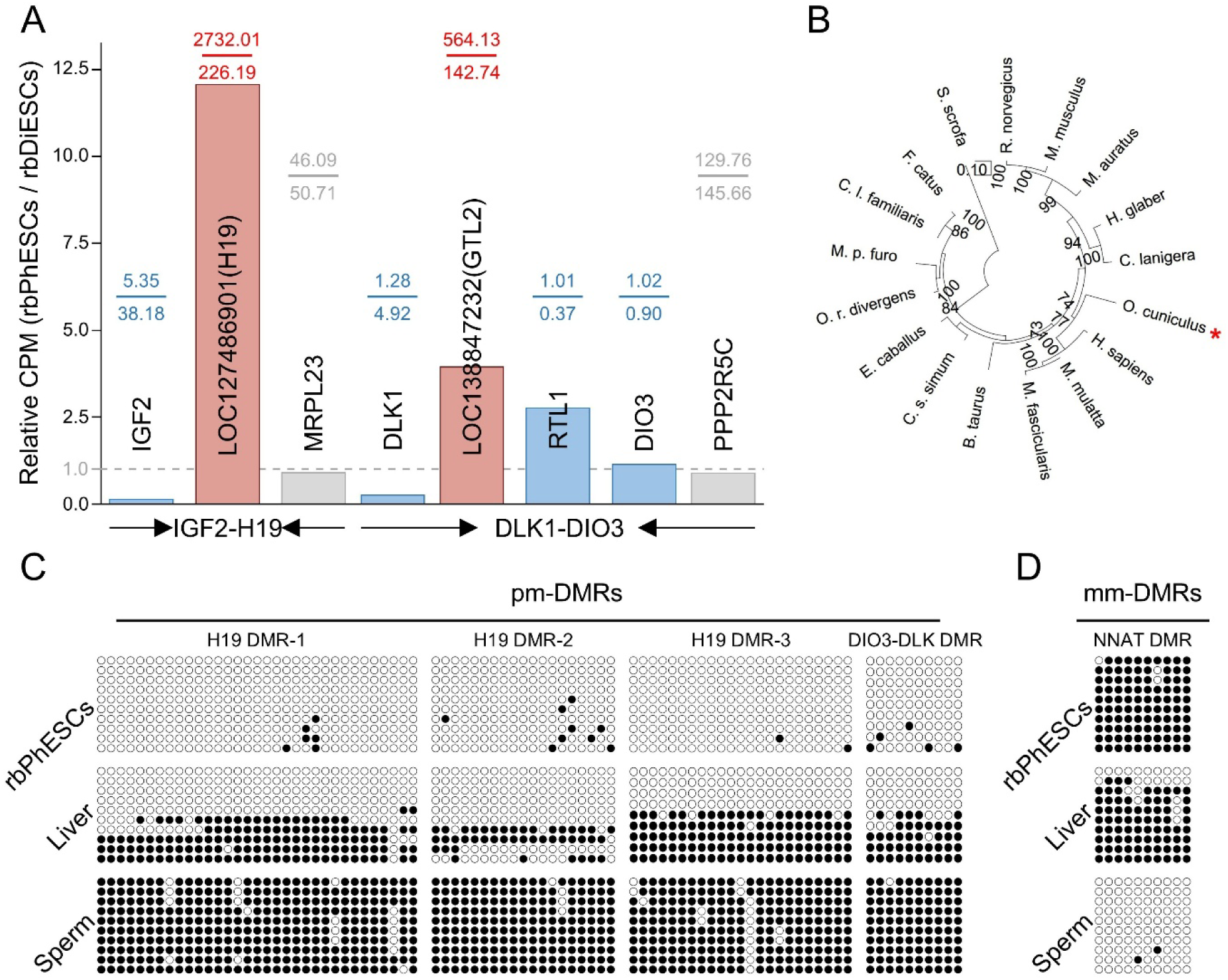
Characterization of Genomic Imprinting in rbPhESCs. (A) Relative expression of genes at the *IGF2*-*H19* and *DLK1*-*DIO3* loci. Expression values were calculated as counts per million (CPM), shown for rbPhESCs relative to rbDiESCs. Red, blue, and grey columns indicate maternal, paternal, and biallelic expression, respectively. Gene symbols are labeled on or above the columns. (B) Evolutionary analysis of *H19* between 17 species. Phylogenetic relationships were inferred using the Maximum Likelihood (ML) method based on the Kimura 2-parameter model of nucleotide substitution. The tree with the highest log likelihood (−20,346.46) is shown. The percentage of replicate trees in which the associated taxa clustered together (1,000 replicates) is shown. The red asterisk marks the position of rabbit (Oryctolagus cuniculus). (**C**) Validation of methylation states at paternally methylated DMRs (pm-DMRs). (**D**) Validation of methylation states at maternally methylated DMRs (mm-DMRs).

Similar to the regulatory mechanism of the *IGF2*-*H19* locus, we found that *GTL2* (annotated as LOC138847232, Gene ID: 138847232), an imprinted gene in *GTL2*-*DIO3* locus, was highly expressed in rbPhESCs compared with rbDiESCs (Fig. 3A), consistent with its maternally expressed pattern. In contrast to previous report, in which *RTL1* and *DIO3* are exclusively expressed in the paternal allele in mice [46], we did not detect them in our fertilized oocyte-derived rbDiESCs (Fig 3A).

Imprinted genes exhibit parent-of-origin–specific monoallelic expression, primarily controlled by DNA methylation in differentially methylated regions (DMRs) [47]. To determine whether imprinting is maintained in rbPhESCs, we performed bisulfite sequencing to compare DNA methylation at classical mammalian DMRs among rbPhESCs, adult male rabbit hepatocytes, and sperm. For *H19*-*IGF2* and *DIO3*-*DLK1* loci, we examined several paternally methylated DMRs (pm-DMRs) and found that most CpG islands were highly methylated in sperm, minimally in rbPhESCs, and intermediately in hepatocytes (Fig. 3C), consistent with those observed in mice [27]. At the DMR of *NNAT*, a gene reported to be maternally methylated [48], rbPhESCs displayed high levels of methylation, while sperm showed little to no methylation, and hepatocytes exhibited intermediate levels (Fig. 3D).

Collectively, these results demonstrate that rbPhESCs retain maternally derived DNA methylation signatures, reflecting their parthenogenetic origin and suggesting their feasibility for use as female gametes.

### Genome-wide genetic screening using rbPhESCs

Because rbPhESCs preserve a stable haploid karyotype even after prolonged culture, we reasoned that they could provide an effective platform for genome-wide forward genetic screening via the one-shot (OS) system [49]. When constructing the OS system, we enhanced the controllability of PiggyBac transposase (PBase) activity by adding two ERT2 domains to both the N- and C-termini of PBase, generating an ERT2–ERT2–PBase–ERT2–ERT2 configuration (Fig. S4A). This design has previously been shown to confer stringent regulation of the nuclear translocation of fused target proteins [50]. We next integrated the advanced OS (aOS) system into the *ROSA26* locus of rbPhESCs (Fig. 4A), thereby generating an rbPhESC line harboring the aOS system (aOS-rbPhESCs) (Fig. 4B, 4C).

**Figure 4.**
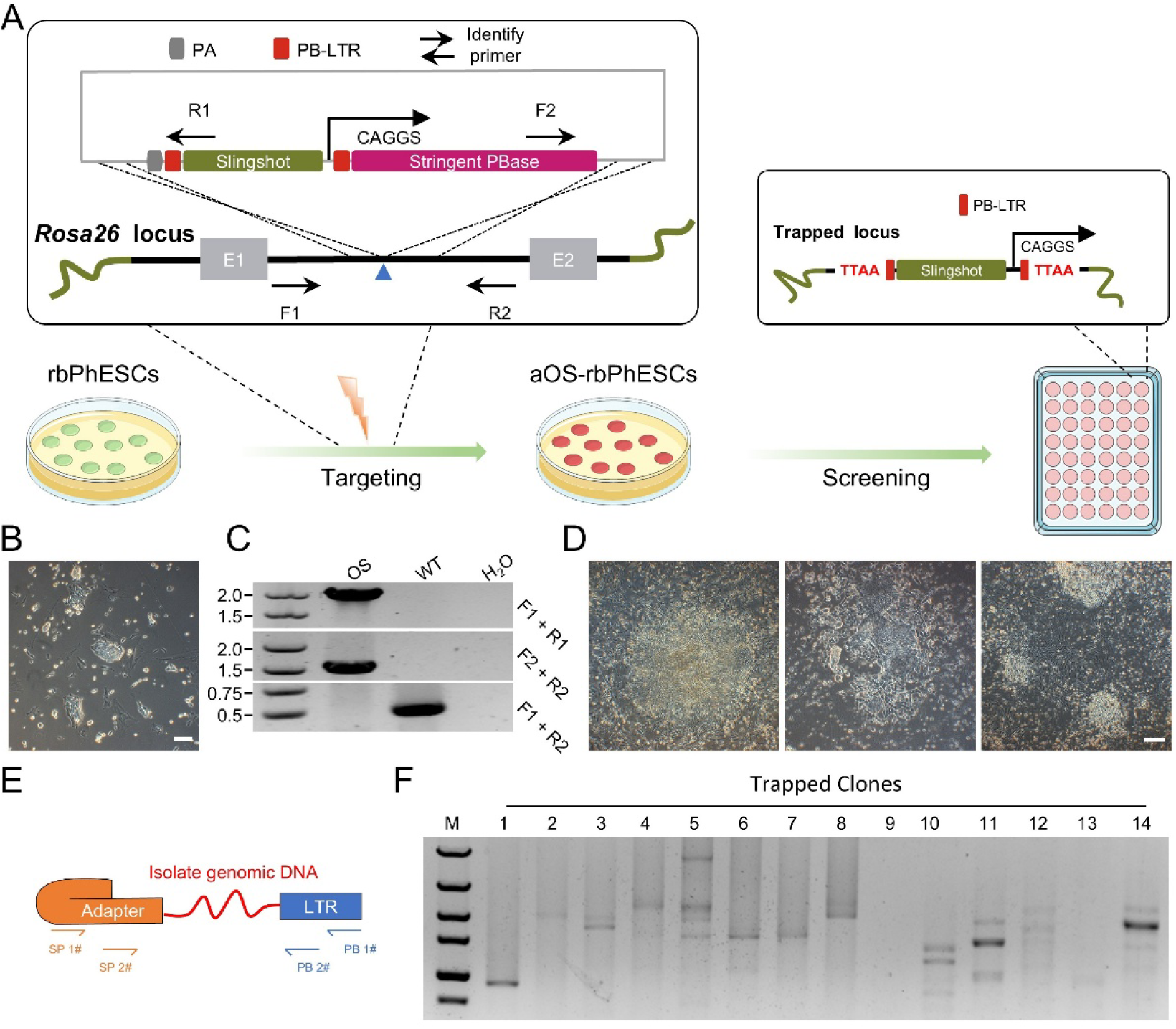
Genome-wide Forward Genetic Screening Using rbPhESCs. **(A)** Schematic of genome-wide screening using the advanced one-shot system in rbPhESCs. An optimized one-shot screening system was established in rbPhESCs by targeting the Slingshot (SA-IRES-Neo-PA) construct into the *ROSA26* locus. To enhance control, a Stringent PBase (ERT2-ERT2-Pbase-ERT2-ERT2 fusion protein) was constructed, allowing tamoxifen-inducible nuclear translocation. Blue triangle: CRISPR-Cas9 cleavage site used to facilitate homology-directed repair (HDR). During the screening phase, the engineered cell population was expanded to 4.5 × 10^7^ cells. Upon treatment with 4-hydroxytamoxifen (4-OHT), PBase translocated into the nucleus, activating genome-wide mutagenesis via the Slingshot trapping cassette. Mutant cells were enriched with G418 selection, and candidate genes conferring BSD resistance were identified by subsequent selection in BSD-containing medium. (**B**) Image of advanced Slingshot-targeted rbPhESCs, referred to as the advanced one-shot system (aOS). Scale bar: 25 µm. (**C**) Gel electrophoresis confirming targeted integration of the Slingshot construct into the *ROSA26* locus in rbPhESCs. (**D**) Representative images of aOS-rbPhESCs during the genome-wide screening process. Left: Cells after 14 days of G418 selection for gene-trapped clones. Middle: Mutant library following 34 days of continued G418 selection. Right: BSD-resistant clones after 7 days of BSD treatment. Scale bar: 25 µm. (**E**) Schematic diagram of Splinkerette PCR. (**F**) Gel electrophoresis image showing trapped loci associated with BSD resistance, identified by Splinkerette PCR.

We next generated a genome-wide mutant library using aOS-rbPhESCs by first enriching the haploid population and expanding it to 4.5 × 10^7^ cells. 4-hydroxytamoxifen (4-OHT) was used to induce nuclear translocation of PBase, which was expected to lead to random, genome-wide insertion of the Slingshot cassette [51]. Insertions that disrupted active loci confer expression of the G418 resistance gene (Fig. S4A), resulting in functionally mutated cells 14 days after G418 selection (Fig. 4D). The yielding mutant library was subsequently subjected to blasticidin (BSD) selection for 7 days. Surviving cells were lysed, and insertion sites were identified by Splinkerette PCR (Fig. 4E, 4F) [52]. These findings demonstrate the successful establishment of a genome-wide mutant library and validate rbPhESCs as a robust system for high-throughput forward genetic screening.

### Generation of maternally depleted *UBE3A* Embryos Using rbPhESCs

Building on our observation that rbPhESCs maintain maternal genomic imprinting patterns, we sought to assess their feasibility for creating animal models of maternal imprinting disorders via semi-cloning approach (Fig. 5A). An eGFP labeled rbPhESC was injected into enucleated metaphase II (MII) oocyte, which was then injected with a sperm into the cytoplasm. Most reconstructed embryos were able to developed into blastocysts expressing eGFP (Fig. 5B), indicating that our rbPhESCs can functionally substitute the maternal genome for making semi-cloning rabbits when combining with sperm injection.

**Figure 5.**
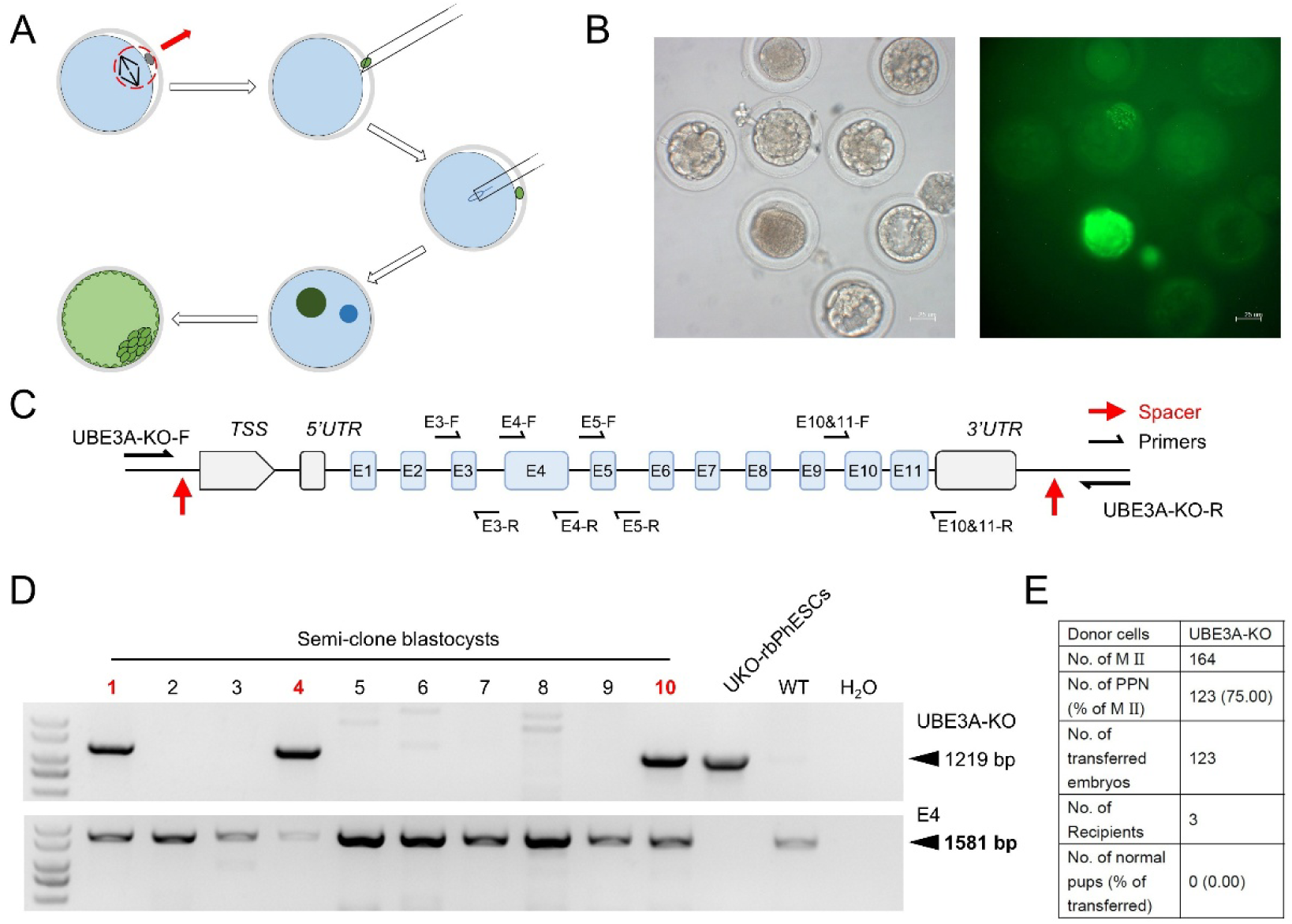
Generation of an imprinting-related disease model using rbPhESCs. (A) Schematic of Semi-Cloning Procedure. Spindle and primary poly body were removed using micropipette, a rbPhESC was then injected into perivitelline space of the enucleated oocyte to serve as a haploid genome donor. Subsequently, a tail-less sperm was introduced assisted by a Piezo-driven micromanipulator. (**B**) Bright-field (left) and fluorescence (right) images of blastocysts derived from semi-cloned rabbit embryos. eGFP-labeled rbPhESCs were used as donor cells. Scale bar: 25 µm. (**C**) Gene structure of rabbit *UBE3A*. Red arrows indicate CRISPR-Cas9 target sites used to delete the entire *UBE3A* locus. Black arrows represent primer positions used for genotyping. (**D**) Gel electrophoresis for identification of *UBE3A* deletion. Primers flanking the UBE3A locus and exon 4 were used for genotyping. Semi-cloned blastocysts showing successful deletion are indicated with red numbers. (**E**) Summary of semi-clone embryos.

We next attempted to explore the feasibility to make rabbit model of Angelman syndrome, which is caused by deficient expression of maternally inherited *UBE3A*. We examined the expression pattern of *UBE3A* and confirmed that the cDNA was truly derived from a single allele (Fig. S5A). To generate a knockout model, we designed sgRNAs targeting sequences flanking the *UBE3A* locus (Table S1) and employed SpCas9 to delete the entire genomic region in rbPhESCs (Fig. 5C). Consequently, we successfully derived three clones with complete *UBE3A* deletion (UKO-1, UKO-4, and UKO-5), in which more than 96 kbps of genomic DNA was excised (Fig. S5B, S5C).

Next, we conducted semi-cloning using the *UBE3A*-knockout rbPhESCs (UKO-4). To generate zygotes carrying UKO-4, chromosomes in MII oocytes were removed by aspirating the meiotic spindle, after which a UKO-4 cell and a tail-less sperm were introduced into the enucleated oocyte (referred to as UKO embryo). Twenty-four hours later, most UKO embryos had reached to the cleavage stage (Fig. S5D), and subsequently developed to the blastocyst stage after an additional three days of culture (Fig. S5E).

To confirm the monoallelic deletion of *UBE3A*, we collected UKO embryos for genotypic analysis. The results showed that the blastocysts carried the same *UBE3A*-deleted allele as UKO-4, along with an intact wild-type allele contributed by the sperm (Fig. 5D). We subsequently transferred 2-cell–stage UKO embryos into the oviducts of surrogate female rabbits. However, no pregnancies were established, and no full-term offspring were obtained (Fig. 5E).

Together, these findings demonstrate that rbPhESCs can serve as maternal genome to support blastocyst development, and may provide a powerful tool for modeling imprinting-related diseases.

## Discussion

In this study, we successfully established rabbit parthenogenetic haploid embryonic stem cells (rbPhESCs) using an optimized culture medium. Consistent with haploid ESCs derived from other mammals [11,53], rbPhESCs harbor a complete haploid chromosome set, exhibit pluripotency comparable to their diploid counterparts, maintain maternal genomic imprinting, and provide a valuable platform for investigating functions of genes. However, our attempts to generate a rabbit model of Angelman Syndrome through semi-cloning—by replacing the maternal genome of reconstructed zygotes with *UBE3A*-deficient rbPhESCs—were unsuccessful.

The haploid nature of haESCs, defined by the absence of allelic redundancy, makes them particularly suitable for efficient studies of gene function. Compared with CRISPR-Cas9-based loss-of-function screens, gene trapping in haploid cells is more random across the genome and independent of sequence context, thereby enabling higher throughput. Leveraging these advantages, haploid cells have been widely applied to identify genes underlying diverse functions, such as toxin receptors [7,54], ESC self-renewal [49,55], and essential genes involved in basic biological processes [56,57].

Consistent with previously described mammalian haESCs, rbPhESCs exhibited enhanced haploid stability, retaining a high proportion of haploid cells for over seven weeks in continuous culture. This prolonged maintenance of haploidy more closely resembles the behavior of monkey haESCs than that of mouse haESCs [9], addressing a major limitation in mammalian haploid stem cell systems and substantially broadens their experimental utility. Moreover, rbPhESCs preserved robust maternal-specific imprinting patterns, with maternal-origin DNA methylation maintained even after extended culture in serum. This stability contrasts with female mouse ESCs, which undergo global hypomethylation at both imprinted and non-imprinted loci when cultured with serum plus LIF, suggesting that rabbit haESCs may possess species-specific mechanisms that better safeguard epigenetic integrity during extended culture [58,59].

In addition, rabbits exhibit an X-chromosome inactivation (XCI) mechanism that closely resembles that of humans but differs markedly from mice. In rabbits and humans, the Xist homologue is not subject to imprinting, and XCI is initiated later than in mice [60]. Furthermore, with appropriate in vitro differentiation strategies, haploidy can be preserved in multiple somatic cell lineages, providing a valuable resource for studying X-chromosome behavior during development [10,54,61]. Collectively, these features make rbPhESCs as a powerful and tractable model for advancing genome imprinting research in a non-rodent mammalian context.

Furthermore, the use of haESCs can greatly enhance both the accuracy and continuity of genome sequence assembly. In conventional genome assembly methods based on diploid cells, substantial effort is required to overcome complications introduced by heterozygosity, segmental duplications, and large structural variations arising from homologous alleles inherited from the mother and father [62]. In contrast, complete hydatidiform mole (CHM) cells, generated by loss of the maternal genome and duplication of the paternal genome post-fertilization, producing a homozygous 46,XX karyotype, were instrumental in assembling the first human telomere-to-telomere (T2T) genome [63]. Similarly, haploid cells, carrying only a single chromosome set and thereby providing a uniform genetic background, have been demonstrated to enable T2T assemblies in mice [20] and crab-eating macaques [21], motivating future efforts to assemble rabbit T2T genomes with our rbPhESCs.

To date, with the exception of mouse and rat models, the generation of complex gene-edited animal models in other species has remained challenging. PhESCs carry chromosomes exclusively of maternal origin and can be used to replace the maternal genome through semi-cloning [26]. Leveraging this property, phESCs provide a distinct advantage for modeling imprinting related diseases, predominantly studied in mice [64], by markedly reducing both the time and complexity required to generate targeted disruptions in parent specific alleles during disease model establishment. This strategy implies that genetic modifications introduced into rbPhESCs could, in principle, be transmitted to offspring if semi-cloning in rabbits can be successfully established in the future, thereby providing a feasible avenue for generating sophisticated rabbit models.

Thus, the establishment of rbPhESCs not only expands the mammalian haploid stem cell repertoire to include rabbits but also holds great promise for broadening the applications of this species in biomedical research.

## Materials and Methods

### Collection of rabbit fertilized embryos

All experiments involving rabbits were approved by the Institutional Animal Care and Use Committees of Guangzhou Institutes of Biomedicine and Health (Animal Welfare Assurance#A5748-01). All donor female rabbits used for embryo collection were sexually mature. Fertilized blastocysts were flushed using flushing buffer (5% FBS in PBS) from the fallopian tubes and uterine horns of donor females 4 days after mating with fertile male rabbits.

To collect fertilized zygotes, donor rabbits were first administered pregnant mare serum gonadotropin (PMSG) at a dose of 30 IU/kg. After 72 hours, PMSG-treated does were mated with fertile males, followed immediately by injection of human chorionic gonadotropin (hCG) at the same dose (30 IU/kg). One-cell stage zygotes were collected from the fallopian tubes 22 hours post-mating and cultured to the blastocyst stage in EBSS complete medium, composed of EBSS (Hyclone, SH30029.02) supplemented with 1× non-essential amino acids (NEAA; Gibco, 11140-050), 1× essential amino acids (EAA; Gibco, 11130-051), 1× GlutaMAX (Gibco, 35050-061), 0.4 mM sodium pyruvate (Gibco, 11360-070), and 10% fetal bovine serum (FBS; Gibco, 10099-141C).

### Collection, activation, and culture of rabbit parthenogenetic embryos

Does in estrus were injected with 30 UI/kg Pregnant Mare Serum Gonadotropin (PMSG, NSHF) intramuscularly to stimulate follicular development. Seventy-two hours after PMSG injection, each doe received an intravenous injection of 30 IU/kg human chorionic gonadotropin (hCG; NSHF) and was immediately mated with vasectomized male rabbits. Thirteen hours later, the superovulated does were sacrificed, and the oviducts (from ovary to uterus) were dissected. Cumulus–oocyte complexes (COCs) were flushed from the oviducts using flushing buffer (5% FBS in PBS) and then, the cumulus cells were removed by treating the COCs with hyaluronidase (3 mg/ml, thermo), followed by gentle pipetting to dissociate the cell layers. After removal of cumulus cells, the oocytes were washed three times in FM (10 % FBS in M199) and transferred to EBSS complete medium.

Mature oocytes without cumulus cells were activated as following steps. Oocytes were activated using electrical pulses (155 V, 10 µs, 3 pulses), followed by incubation for 5 minutes in Ionomycin medium, consisting of EBSS complete medium supplemented with 10 µM Ionomycin (Sigma, I0634-1mg). The oocytes were then cultured for 1 hour in 6-DX medium, prepared with EBSS complete medium supplemented with 2 mM 6-DMAP (Sigma, D2629-100MG) and 5 µg/ml cycloheximide (CHX; Sigma, 46401-100MG-R). Activated embryos were subsequently cultured in EBSS complete medium at 38.5 °C in an atmosphere of 5% CO_2_ in air for an additional 4 days, until the blastocyst appeared apparently.

### Derivation of Rabbit Haploid Embryonic Stem Cells (rbPhESCs) from Parthenogenetic Embryos

Blastocysts derived from parthenogenetically activated embryos were selected and transferred to 4-well dishes pre-coated with feeder cells and maintained in iKFC medium, consisting of DMEM/F-12 (Gibco, 11330-032) supplemented with 10% KnockOut™ Serum Replacement (Gibco, 10828028), 10% fetal bovine serum (Gibco, 10099-141c), 1× GlutaMAX (Gibco, 35050-061), 1× non-essential amino acids (Gibco, 11140-050), 1× penicillin–streptomycin (Gibco, 15140-163), 50 μM β-mercaptoethanol (Thermo Fisher, 21985023), 8 ng/mL recombinant human FGF-basic (PeproTech, 100-18B-100), 1 μM CHIR99021 (Selleck, S1263), and 1 μM IWR-endo-1 (Selleck, S7086). Approximately 7 days after seeding, outgrowths appeared adjacent to the blastocysts. These outgrowths were manually picked, dissociated into single cells, and replated onto new feeder-coated 4-well dishes. Emerging colonies were subsequently passaged multiple times using 0.05% trypsin until a sufficient cell population was obtained for fluorescence-activated cell sorting (FACS).

### Enrichment of Haploid rbPhESCs by FACS

To enrich the haploid population, exponentially proliferating rbPhESCs were dissociated into single cells using 0.05% trypsin and stained with Hoechst 33342 (5 µg/mL) at 37 °C for 15 minutes. During staining, the cell suspension was gently agitated every 5 minutes to ensure uniform staining. After incubation, cells were filtered through a 40 µm strainer to remove clumps and ensure a single-cell suspension. The DNA content of individual cells was analyzed by flow cytometry (Beckman Coulter MoFlo AstriosEQ Cell Sorters), and haploid cells (1n population) were identified and sorted. Sorted cells were then seeded onto feeder-coated 4-well dishes and cultured in iKFC medium.

### Karyotyping analysis

Cells at approximately 80% confluence were treated with 50 ng/mL colcemid for 2 hours to arrest cells in metaphase. After treatment, cells were dissociated and incubated at 37 °C for 30 minutes in a hypotonic solution containing 0.075 M potassium chloride (Sigma, P5405) in PBS (Gibco, 10010-023). Following hypotonic treatment, cells were fixed with at least three changes of freshly prepared fixative composed of methanol (Sigma, 34860) and acetic acid (Sigma, 695092) in a 3:1 ratio. Metaphase chromosome spreads were prepared by dropping the fixed cells onto clean glass slides and stained using a standard G-banding protocol.

### De Novo Genome Sequencing and Copy Number Variation (CNV) Analysis

Cells in the exponential growth phase were dissociated and used for genomic DNA extraction. Freshly extracted genomic DNA was used to prepare sequencing libraries following the standard workflow for the DNBSEQ platform, using the MGI Easy Universal DNA Library Prep Kit (MGI, 1000006985). After sequencing, raw reads were filtered using fastp [65] with the following parameters: fastp -w 48 -z 4 -q 20 -u 30 -n 10. Filtered reads were aligned to the rabbit reference genome (mOryCun1.1, GCF_964237555.1) using BWA with default parameters [66]. Duplicate alignments were removed using sambamba [67] with the following command: sambamba markdup -r -l 5. For whole-genome CNV analysis, the genome was divided into 100,000 bp bins, and FPKM-normalized read coverage was calculated using deepTools [68] with the following parameters: bamCoverage --binSize 100000, --numberOfProcessors, ‘max’ -- effectiveGenomeSize 2804657419 --normalizeUsing ‘RPKM’ --outFileFormat ‘bedgraph’. For exon-level variation analysis, the genome was divided into 1,000 bp bins, and coverage was calculated with the same tool and similar parameters, adjusted for bin size: bamCoverage -- binSize 1000, --numberOfProcessors, ‘max’ --effectiveGenomeSize 2804657419 -- normalizeUsing ‘RPKM’ --outFileFormat ‘bedgraph’. Fold-change in CNV was calculated using the formula: FC(CNV) = FPKM(haploid bin coverage) / FPKM(diploid bin coverage). Bins with FC(CNV) > 2 were identified and compared to annotated exon regions in the rabbit reference genome to determine overlap.

### Immunofluorescent staining of rbPhESCs

Immunofluorescent staining was performed using a standard protocol. Briefly, rbPhESCs were seeded onto feeder-coated round glass coverslips placed in 4-well dishes. After four days of culture, the coverslips were transferred to new 4-well dishes and fixed with 4% paraformaldehyde (Sigma, 158127). Following three washes with PBS, the cells were permeabilized and incubated with primary antibodies against *OCT4* (Santa Cruz Biotechnology, sc-5279), *SOX2* (R&D, MAB2018), and *NANOG* (R&D, AF1977). After additional washing, appropriate secondary antibodies (Abcam, AlexaFluor) were applied. Fluorescence images were then acquired using a confocal laser scanning microscope (Zeiss 710 NLO).

### Alkaline phosphatase (AP) staining

Cells were seeded and cultured in a 6-well plate until they reached the exponential growth phase, then fixed with 4% paraformaldehyde (PFA). Alkaline phosphatase (AP) staining was performed according to the manufacturer’s instructions (Beyotime, C3206), and stained cells were imaged using an upright optical microscope.

### Embryoid Body Formation

Exponentially expanded rbPhESCs were dissociated and washed three times with PBS, then resuspended in iKFC medium lacking CHIR99021, IWR-1, and KOSR. The cell suspension was plated in 10 cm low-attachment culture dishes and incubated for 14 days. Images were captured on day 3 and day 8 using an inverted optical microscope.

### Teratoma Formation and Histological Analysis

A total of 5 × 10^7^ exponentially proliferating rbPhESCs were collected and resuspended in 0.3 mL of 0.9% sodium chloride solution. Ten-week-old male severe combined immunodeficiency (SCID) mice were injected subcutaneously with the cell suspension and housed in a specific pathogen-free (SPF) facility under controlled conditions (22 ± 1 °C, 50 ± 10% humidity, 12-hour light/dark cycle). Eight weeks post-injection, teratomas were harvested and processed for histological examination using standard hematoxylin and eosin (H&E) staining.

### RNA-Sequencing and Data Processing

Total RNA was extracted from each sample and reverse-transcribed into cDNA using the SuperScript™ II Reverse Transcriptase (Invitrogen, 1896649, USA). Purified cDNA was then employed for sequencing library preparation with the Illumina Nextera XT DNA Library Preparation Kit (Illumina, FC-131-1001), according to the manufacturer’ s instructions. Raw RNA-Seq reads were filtered using fastp [65] with the following parameters: fastp -w 48 -z 4 -q 20 -u 30 -n 10. Filtered reads were aligned to rabbit reference genome (mOryCun1.1, GCF_964237555.1) using HISAT2 [69] with the command: hisat2 -x hs2IdxBase --known-splicesite-infile pathRefGenomeSS --novel-splicesite-outfile pathSampleS -p 24. Potential duplicate alignments were removed using sambamba [67] with the following command: sambamba markdup -r -t 24 -l 5.

Cleaned BAM files were further analyzed using the R packages Rsubread [70] and edgeR [71]. Briefly, gene-level read counts were calculated using the featureCounts with the following parameters: featureCounts(files = bamPath, annot.ext = pathGtf, isGTFAnnotationFile = T, GTF.featureType = “exon”, GTF.attrType = “gene_id”, isPairedEnd = TRUE, nthreads = 24). Lowly expressed genes were filtered using filterByExpr() in edgeR, and normalized using counts per million (CPM) via the cpm() function.

Hierarchical clustering heatmaps for global transcriptomes and lineage-specific genes were generated using the ComplexHeatmap package [72]. Scatter plots were produced with ggplot2 [73], and pathway and Gene Ontology (GO) enrichment analyses were performed using DAVID [74,75].

### Evolutionary Relationships Analysis of mammal *H19*

DNA sequences of the *H19* gene from 17 species were aligned using MUSCLE in MEGA12 [76] with default parameters. Phylogenetic analysis was then performed using the Maximum Likelihood (ML) method under the Kimura 2-parameter model of nucleotide substitution [77]. The resulting tree with the highest log-likelihood score (−20,346.46) is shown, with node support evaluated using 1,000 bootstrap replicates [78]. For the heuristic search, the initial tree was chosen based on the higher log-likelihood between a Neighbor-Joining (NJ) tree [79] and a Maximum Parsimony (MP) tree. The NJ tree was constructed from a pairwise distance matrix calculated using the Kimura 2-parameter model [77], while the MP tree was the shortest among 10 trees generated from random starting points. The final dataset included 17 nucleotide sequences with 2,936 aligned positions. All analyses were conducted in MEGA12 [76], utilizing up to seven parallel computing threads.

### Detection of DNA methylation in DMR

A total of 2 μg of genomic DNA from each sample was subjected to bisulfite conversion using either the EpiTect Fast DNA Bisulfite Kit (QIAGEN, 59824) or the EpiArt Ultrafast DNA Methylation Bisulfite Kit (Vazyme, EM112-01), following the manufacturer’s instructions. The DMRs of *H19*, *IG*, and *NNAT* were amplified by PCR using KOD One™ PCR Master Mix BLUE (TOYOBO, KMM-201NV) under the conditions of 95 °C for 3 min; 45 cycles of 95 °C for 10 s, 55 °C for 30 s, 68 °C for 10 s; followed by a final extension at 68 °C for 5 min and a hold at 12 °C. PCR products were cloned into the pMD18-T vector using the Hieff Clone® Universal Zero TOPO TA/Blunt Cloning Kit (Yeasen, 10906ES) and sequenced using the M13 forward primer. All primers used for fragment amplification are listed in Table S2.

### Construction of Advanced One-shot (aOS) Plasmids

The PiggyBac transposon vector (Systembio, PB513B-1) and PiggyBac transposase (Systembio, PB220PA-1) were obtained from System Biosciences, LLC. The ERT2–ERT2 tandem domains were synthesized by a commercial provider, and the PBase coding sequence was cloned from PB220PA-1. To enable fusion of PBase with the ERT2–ERT2 domains, linker peptides (GRPESGA at the N-terminus and SRADPKKKRKVDI at the C-terminus) were inserted between PBase and ERT2. The resulting ERT2-fused PBase construct, along with the Slingshot cassette, was cloned into the PB513B-1 backbone to generate the final aOS plasmid (Supplementary sequences 2).

### Generation of aOS-rbPhESC Screening System

rbPhESCs were dissociated and co-electroporated with the px330-xx plasmid carrying sgRNAs targeting the genomic region between exon 1 and exon 2 of the rabbit *ROSA26* locus (sgRNA sequences listed in Table S1), along with the aOS plasmid. Stably integrated clones were selected using 500 ng/mL puromycin (MedchemExpress, HY-B1743) and verified by PCR using primers listed in Table S2.

### Splinkerette PCR for Isolation of Insertion Sites

Genomic DNA was extracted from cells harboring Slingshot cassette insertions. One microgram of genomic DNA was digested with Sau3A I and ligated to a Splinkerette adaptor. One microliter of the ligation product was used as the template for two rounds of nested PCR. Adaptor sequences and primers are detailed in Table S2.

### Semi-clone and Embryo Transfer

rbPhESCs within two passages post-FACS sorting were used as donor cells for semi-cloning. Metaphase II (M II) oocytes were collected as described in the rbPhESC derivation protocol and placed in M199 medium (Gibco, 12350-039) supplemented with 7.5 μg/mL cytochalasin B (Sigma, C6762-5MG). After 10 minutes of incubation, the polar body and meiotic spindle were removed using a micromanipulator with a sharp glass needle. A single rbPhESC was then injected into the perivitelline space. Freshly collected spermatozoa were subjected to ultrasonic treatment to remove the tails, and a tail-less sperm was injected into the cytoplasm of the enucleated oocyte using a blunt needle assisted by a Piezo-driven micromanipulator. Reconstructed oocytes were then fused, activated, and cultured as previously described. Embryos that reached the 2-cell stage were transferred into the oviducts of surrogate females that had been mated with vasectomized males 48 hours prior to embryo transfer [80].

### Reverse Transcription PCR

Freshly isolated tissues were manually homogenized and lysed using Trizol regent (Invitrogen, 15596026) following the manufacturer’s instructions. Total RNA was extracted and reverse transcribed into complementary DNA (cDNA) using the random primers (Promega, C118), M-MLV Reverse Transcriptase (Promega, M1701), Recombinant RNasin Ribonuclease Inhibitor (Promega, N251) and a dNTP mix containing dATP, dCTP, dGTP, and dTTP (Promega, U1330). Both cDNA and genomic DNA were used as templates for PCR, with primers listed in Table S2. PCR products were subsequently sequenced and analyzed to determine the allelic origin of the cDNA.

## Supporting information

Supplementary sequences

## Acknowledgements

We thank Minghui Gao for the help with FACS. This work was supported by the National Natural Science Foundation of China (32400423, 32100409), the Science and Technology Planning Project of Guangdong Province (2022A1515012058), the Science and Technology Program of Guangzhou (202201010617), Major Research Project of Guangzhou Institutes of Biomedicine and Health, Chinese Academy of Sciences (GIBHMRP25-01), Science and Technology Program of Guangzhou, China (2024B03J1231).

