## Supplementary Figures for "Derivation, Characterizations, and Applications of Rabbit Haploid Embryonic Stem Cells"

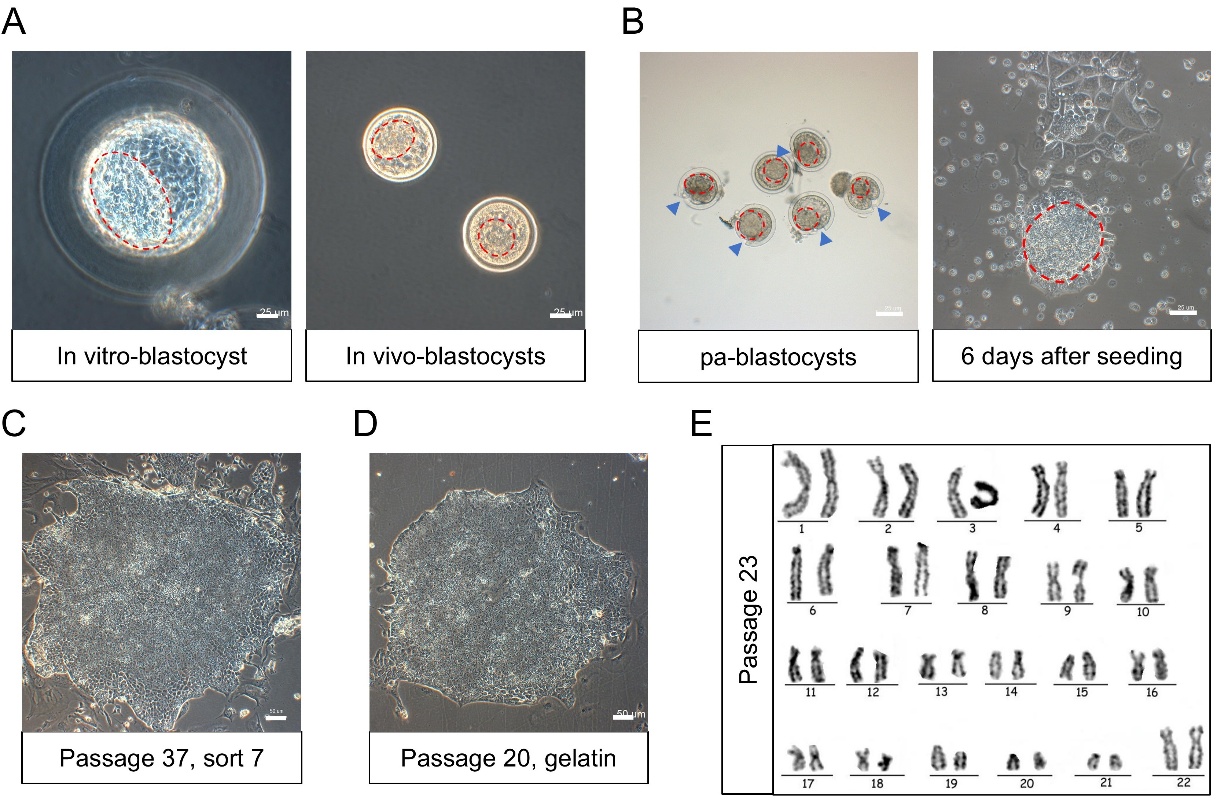

**Supplementary Figure 1 | Derivation of rbPhESCs from in vitro–activated rabbit oocytes**

(**A**) Bright-field images of rabbit in vivo–developed (left) and in vitro–developed embryos (right). Inner cell masses (ICMs) within the blastocoels are outlined with red dashed lines. Fertilized blastocysts (left) were recovered from the fallopian tubes 96 hours after mating with a male rabbit, whereas zygotes were flushed 20 hours post-mating and subsequently cultured for 76 hours after collection (right). Scale bar: 25 µm. (**B**) Representative images of in vitro–developed parthenogenetic blastocysts (left) and their outgrowths (right). Inner cell masses (ICMs) within the blastocoels are outlined with red dashed lines; mechanically introduced nicked lesions on the zona pellucida are indicated with blue triangles; compact outgrowths are outlined with red dashed line. pa-blastocysts: parthenogenetic blastocysts. Scale bar: 25 µm. (**C**) Phase-contrast image of rbPhESCs after continuous culture for approximately four months. Scale bar: 50 µm. (**D**) Colony morphology of rbPhESCs cultured on gelatin for five days, scale bar: 50 µm. (**E**) Karyotype analysis of rabbit fertilized diploid ESCs, numbers under the chromosomes are randomly assigned, not represent the number of chromosomes.

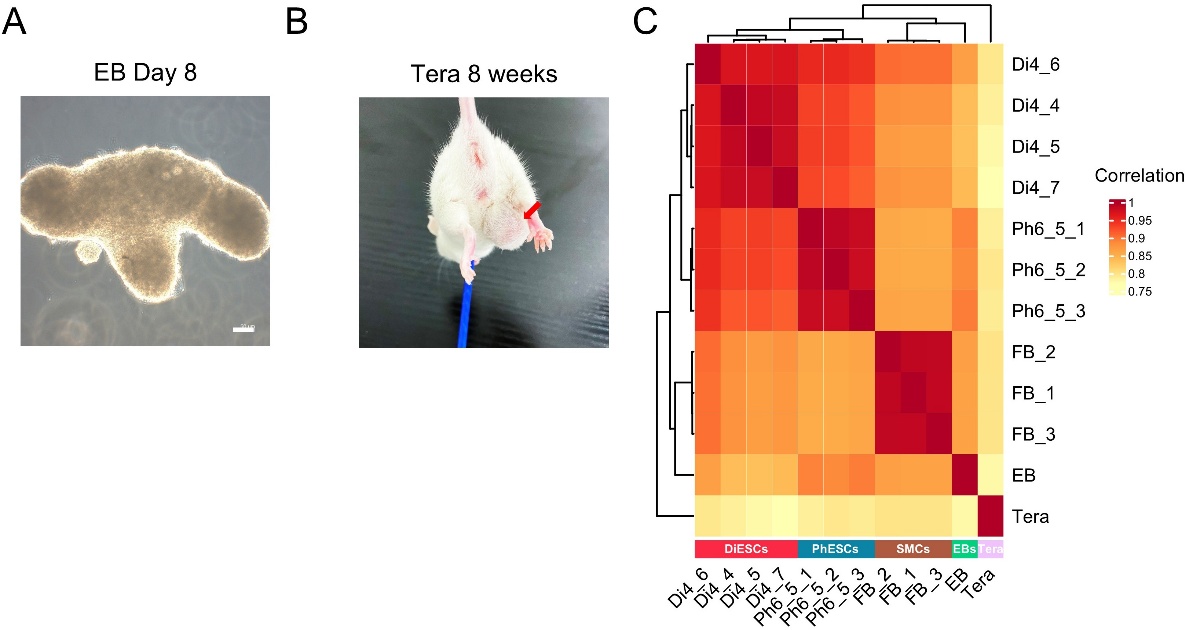

**Supplementary figure 2 | Pluripotency Characterization of rbPhESCs**

(**A**) Embryoid body after 8 days of differentiation in iKFC medium without CHIR 99021 and IWR-1. Scale bar: 50 µm. (**B**) Teratoma formation in SCID mice. Approximately 2 × 10^6^ rbPhESCs were dissociated, suspended in physiological saline, and injected into the inguinal region of severe combined immunodeficiency (SCID) mice. Images of tumors were captured at the injection site 8 weeks post-injection. (**C**) Hierarchical clustering of samples based on expressed genes with counts per million (CPM). Lowly expressed genes were filtered using edgeR. Di: rabbit diploid embryonic stem cells; Ph: rabbit parthenogenetic embryonic stem cells; EB: embryonic body; Tera: teratocarcinomas; FB: fibroblasts.

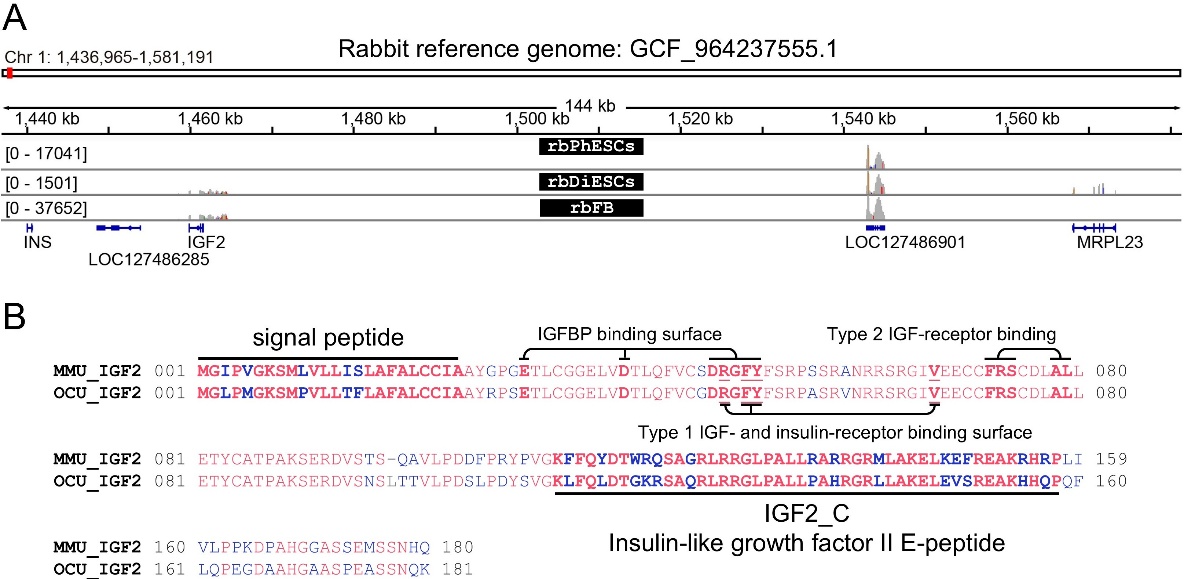

**Supplementary figure 3 | Genes located in *IGF2*-*H19* locus.**

(**A**) RNA-Seq read distribution across the *IGF2*–*H19* imprinted locus spanning the genomic region between *INS* and *MRPL23*. (**B**)Comparison of *IGF2* amino acid sequences between mouse (upper line) and rabbit (lower line). Functional domains are indicated by corresponding labels. Amino acid substitutions are highlighted in blue, and insertions/deletions (indels) are shown in gray.

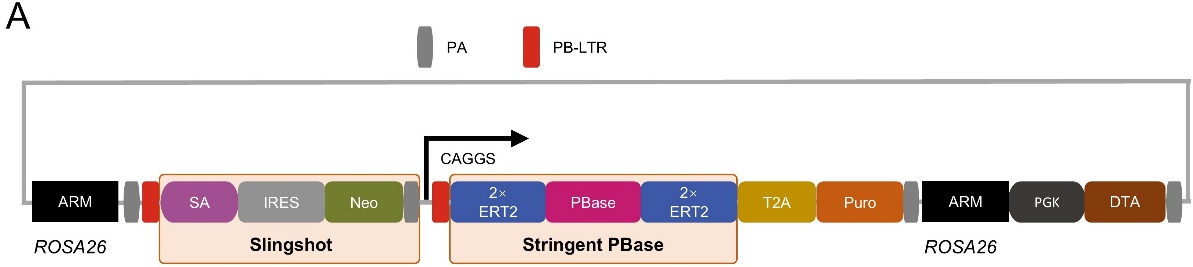

**Supplementary figure 4 | Schematic diagram of the advanced one-shot (aOS) system.**

(**A**) A diagram illustrating the design of the aOS construct, which includes the inducible PBase flanked by tandem ERT2 domains at both termini and the Slingshot gene-trapping cassette. The entire system is flanked by PiggyBac long terminal repeats (PB-LTRs) and targeted into the *ROSA26* locus of rbPhESCs for controlled, genome-wide mutagenesis upon 4-OHT induction.

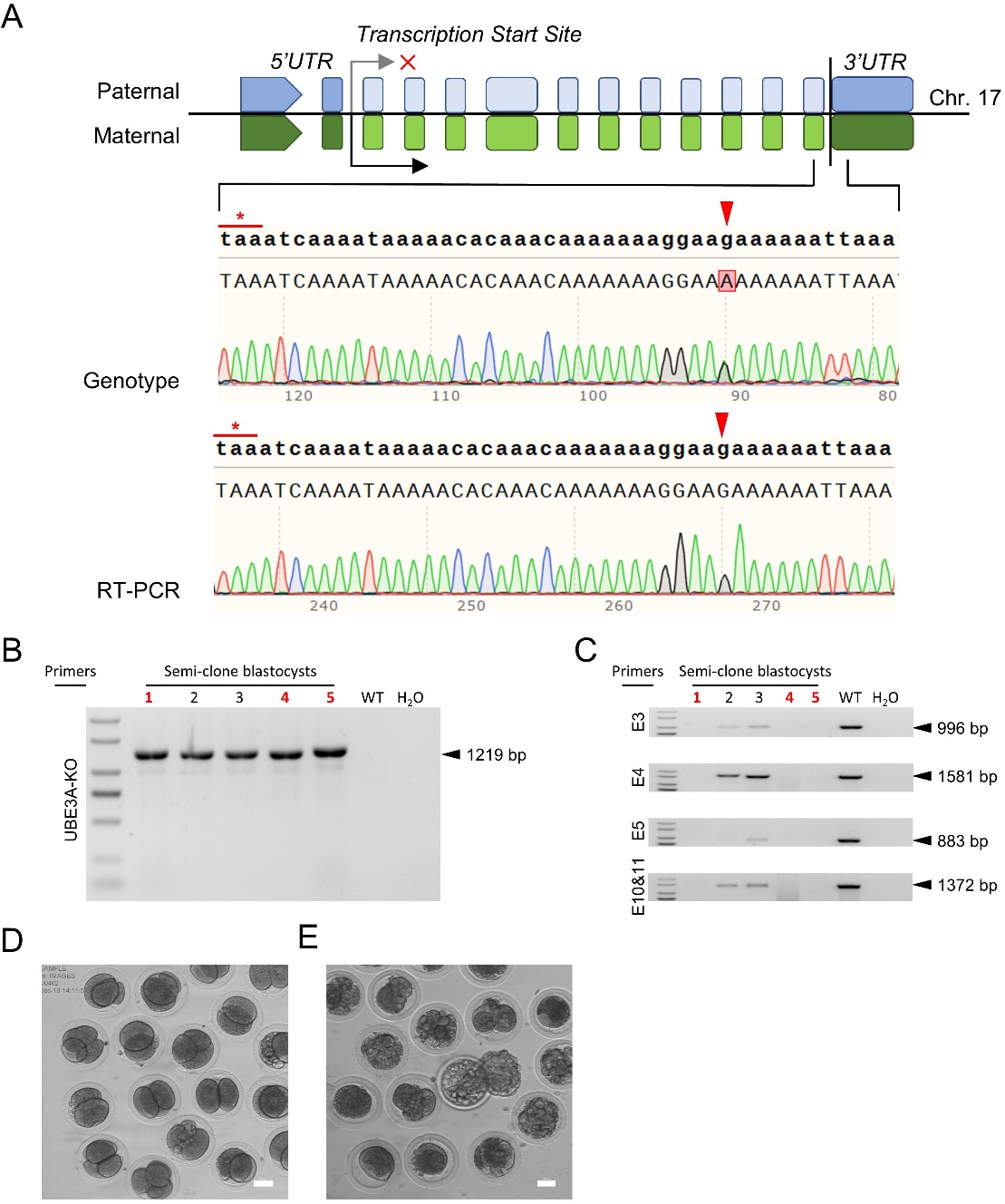

**Supplementary figure 5 | Generation of maternal depletion of *UBE3A* model using rbPhESCs.**

(**A**) Imprinting pattern and expression validation of *UBE3A* in rabbit olfactory bulb. (**B**) Gel electrophoresis for identification of *UBE3A* deletion. Primers flanking the *UBE3A* locus were used for genotyping. Clones of rbPhESCs showing successful deletion are indicated with red numbers. (**C**) Gel electrophoresis of PCR products targeting internal *UBE3A* exons to confirm sequence deletion. Positive cell clones are marked with red numbers. (**D**) 2-cell semi-cloned embryos generated using rbPhESCs as maternal genome and sperm as paternal genome. Scale bar: 50 µm. (**E**) Blastocysts of semi-cloned embryos developed from panel D. Scale bar: 50 µm.
