## Supplementary Tables for "Derivation, Characterizations, and Applications of Rabbit Haploid Embryonic Stem Cells"

Supplementary information, Table S1 sgRNA used in the study

| Application | Forward | Reverse |
| --- | --- | --- |
| *ROSA26* targeting sgRNA | CACCGACTCGGGCCTCATGGTCTCGAGG | AAACCCTCGAGACCATGAGGCCCGAGTC |
| *UBE3A* knock out sgRNA1 | CACCGATAGGCCGCGAGCTACTCCG | AAACCGGAGTAGCTCGCGGCCTATC |
| *UBE3A* knock out sgRNA2 | CACCGTCTCGCGACCGCGCGCAAGA | AAACTCTTGCGCGCGGTCGCGAGAC |
| *UBE3A* knock out sgRNA3 | CACCGACTGCTATCGAATCAGTGGA | AAACTCCACTGATTCGATAGCAGTC |
| *UBE3A* knock out sgRNA4 | CACCGTACTGTGATAAGGTAACGTG | AAACCACGTTACCTTATCACAGTAC |

Supplementary information, Table S2 Primers used in the study

| Application | Primer | Sequence | Product size (bp) | Annealing (℃) |
| --- | --- | --- | --- | --- |
| Recombination arm amplification | *ROSA26*-short-F | GAAGGGCGATCTCAACCTGCACAGTCAAGT | 554 | 60 |
|  | *ROSA26*-short-R | TGTTTAAACGCCTGTTCTAACCATCACTGT |  |  |
|  | *ROSA26*-long-F | CTCATCAATGTATCTTATCACGCGTTCGAGACCATGAGGCCCGAG | 770 | 52 |
|  | *ROSA26*-long-R | GATGGCGCGCCACAATGCCCATGCCTTCAAC |  |  |
| *ROSA26* genotyping | *ROSA26*-WT-F | TGCTAGGCCTGTTTTCCTGA | 725 | 55 |
|  | *ROSA26*-WT-R | GGAGACCTCTCTCTCTCTGC |  |  |
|  | *ROSA26*-OS-5ARM-F | GGGTGGAGCCAGGAACTTTT | 1467 | 58 |
|  | *ROSA26*-OS-5ARM-R | CGCAACCTCCCCTTCTACGA |  |  |
|  | *ROSA26*-OS-3ARM-F | CCTCCACGGGTTCAAAAACG | 1674 | 58 |
|  | *ROSA26*-OS-3ARM-R | AAAAGGTGAGAAACAGGCAGA |  |  |
| *UBE3A* genotyping | *UBE3A*-WT-E3-F | ATCTGTGTTAAGTGCAAATCAAGT | 996 | 55 |
|  | *UBE3A-*WT-E3-R | GCTTGGAGTTTAGGATTGGTAGC |  |  |
|  | *UBE3A*-WT-E4-F | TGTTGAGGCTGTTGATATTGGTA | 1581 | 55 |
|  | *UEB3A*-WT-E4-R | AATACCCGTGTGTTCCTGTGA |  |  |
|  | *UBE3A*-WT-E5-F | TGTTTGGGTTCTTTCCATGTTGA | 883 | 57 |
|  | *UEB3A*-WT-E5-R | ACTCCCCAAGAAACCTCATAGTG |  |  |
|  | *UBE3A*-Exon10&11-F | AGGGAGTTCTGGGAAATCGT | 1372 | 56 |
|  | *UBE3A*-Exon10&11-R | GAGGCACAGACAGAGGTGAC |  |  |
|  | *UBE3A*-KO-F | GCTCTTCTGTTGGGCCGATT | 1219 | 58 |
|  | *UBE3A*-KO-R | CTTCCTGTAGACTGTGGGCTG |  |  |
| *UBE3A* RT-PCR | *UBE3A*-Exon10&11-F | AGGGAGTTCTGGGAAATCGT | 411 | 58 |
|  | *UBE3A*-Exon10&11-R | GAGGCACAGACAGAGGTGAC |  |  |
| Bisulite sequencing of DMRs | *H19*-DMR1-out-F | GAATTGGGGAGTAGGGAGAT | 510 | 55 |
|  | *H19*-DMR1-out-R | CCTAATACTCCCCAACTTCT |  |  |
|  | *H19*-DMR1-in-F | TGTTGTTTAGGAGTAAAAGT | 482 | 52 |
|  | *H19*-DMR1-in-R | TCAACTTCCCCAATACAACT |  |  |
|  | *H19*-DMR2-out-F | ATATTTGGGTGTGGGATAAA | 431 | 55 |
|  | *H19*-DMR2-out-R | CCAACATCCTTTACCATCTC |  |  |
|  | *H19*-DMR2-in-F | AGGGGATTTGAGTTTAGGGT | 375 | 52 |
|  | *H19*-DMR2-in-R | CTTCCCCAATACAACTCAAA |  |  |
|  | *H19*-DMR3-out-F | TTTGAGTTGTATTGGGGAAG | 319 | 55 |
|  | *H19*-DMR3-out&in-R | CTAAAATACAAAAAAACCCCA |  |  |
|  | *H19*-DMR3-in-F | ATGTTTTGTGTGATGGTGAT | 257 | 52 |
|  | *H19*-DMR3-out&in-R | CTAAAATACAAAAAAACCCCA |  |  |
|  | *IG*-DMR-F | TATTTTAGGGTGTTTGGAGG | 148 | 50 |
|  | *IG*-DMR-R | AATACCCTAAAATTCAAACC |  |  |
|  | *NNAT*-DMR-out-F | GGTAGAGGTTGAAAGGATTTGG | 476 | 54 |
|  | *NNAT*-DMR-out-R | CCCCTTCCAAAAAATTCCGCCT |  |  |
|  | *NNAT*-DMR-in-F | GGGATTTTTGGGTAGTAGAGAATT | 185 | 54 |
|  | *NNAT*-DMR-in-R | AATACCCCTCTTTCTAAACCCTAAC |  |  |
| Splinkerette-PCR for one-shotting | Long-strand adaptor | CGAAGAGTAACCGTTGCTAGGAGAGACCGTGGCTG-AATGAGACTGGTGTCGACACTAGTGG | _ | _ |
|  | Short-strand adaptor | GATCCCACTAGTGTCGACACCAGTCTCTAATTTTTTTT-TTCAAAAAAA | _ | _ |
|  | Splink1 | CGAAGAGTAACCGTTGCTAGGAGAGACC | _ | 58 |
|  | 1st-U3 LTR-R | AGACCGATAAAACACATGCGTC |  |  |
|  | Splink2 | GTGGCTGAATGAGACTGGTGTCGAC | _ | 58 |
|  | 2nd-U3 LTR-R | CGCATGATTATCTTTAACGTACGTC |  |  |
