## Supplementary sequences for "Derivation, Characterizations, and Applications of Rabbit Haploid Embryonic Stem Cells": Supplementary sequence 2-ERT2 fused PBase.docx

**Supplementary sequences 2 | ERT2 fused PBase**

ERT2

Linker-1

Linker-2

PBase

GCTGGAGACATGAGAGCTGCCAACCTTTGGCCAAGCCCGCTCATGATCAAACGCTCTAAGAAGAACAGCCTGGCCTTGTCCCTGACGGCCGACCAGATGGTCAGTGCCTTGTTGGATGCTGAGCCCCCCATACTCTATTCCGAGTATGATCCTACCAGACCCTTCAGTGAAGCTTCGATGATGGGCTTACTTACCAACCTGGCAGACAGGGAGCTGGTTCACATGATCAACTGGGCGAAGAGGGTGCCAGGCTTTGTGGATTTGACCCTCCATGATCAGGTCCACCTTCTAGAATGTGCCTGGCTAGAGATCCTGATGATTGGTCTCGTCTGGCGCTCCATGGAGCACCCAGTGAAGCTACTGTTTGCTCCTAACTTGCTCTTGGACAGGAACCAGGGAAAATGTGTAGAGGGCATGGTGGAGATCTTCGACATGCTGCTGGCTACATCATCTCGGTTCCGCATGATGAATCTGCAGGGAGAGGAGTTTGTGTGCCTCAAATCTATTATTTTGCTTAATTCTGGAGTGTACACATTTCTGTCCAGCACCCTGAAGTCTCTGGAAGAGAAGGACCATATCCACCGAGTCCTGGACAAGATCACAGACACTTTGATCCACCTGATGGCCAAGGCAGGCCTGACCCTGCAGCAGCAGCACCAGCGGCTGGCCCAGCTCCTCCTCATCCTCTCCCACATCAGGCACATGAGTAACAAAGGCATGGAGCATCTGTACAGCATGAAGTGCAAGAACGTGGTGCCCCTCTATGACCTGCTGCTGGAGGCGGCGGACGCCCACCGCCTACATGCGCCCACTAGCCGTGGAGGGGCATCCGTGGAGGAGACCGACCAAAGCCACTTGGCCACTGCGGGCTCTACTTCATCGCATTCCTTGCAAAAGTATTACATCACGGGGGAGGCAGAGGGTTTCCCTGCCACAGCTCCTAGGGGAGGCTCTGCTGGAGACATGAGAGCTGCCAACCTTTGGCCAAGCCCGCTCATGATCAAACGCTCTAAGAAGAACAGCCTGGCCTTGTCCCTGACGGCCGACCAGATGGTCAGTGCCTTGTTGGATGCTGAGCCCCCCATACTCTATTCCGAGTATGATCCTACCAGACCCTTCAGTGAAGCTTCGATGATGGGCTTACTTACCAACCTGGCAGACAGGGAGCTGGTTCACATGATCAACTGGGCGAAGAGGGTGCCAGGCTTTGTGGATTTGACCCTCCATGATCAGGTCCACCTTCTAGAATGTGCCTGGCTAGAGATCCTGATGATTGGTCTCGTCTGGCGCTCCATGGAGCACCCAGTGAAGCTACTGTTTGCTCCTAACTTGCTCTTGGACAGGAACCAGGGAAAATGTGTAGAGGGCATGGTGGAGATCTTCGACATGCTGCTGGCTACATCATCTCGGTTCCGCATGATGAATCTGCAGGGAGAGGAGTTTGTGTGCCTCAAATCTATTATTTTGCTTAATTCTGGAGTGTACACATTTCTGTCCAGCACCCTGAAGTCTCTGGAAGAGAAGGACCATATCCACCGAGTCCTGGACAAGATCACAGACACTTTGATCCACCTGATGGCCAAGGCAGGCCTGACCCTGCAGCAGCAGCACCAGCGGCTGGCCCAGCTCCTCCTCATCCTCTCCCACATCAGGCACATGAGTAACAAAGGCATGGAGCATCTGTACAGCATGAAGTGCAAGAACGTGGTGCCCCTCTATGACCTGCTGCTGGAGGCGGCGGACGCCCACCGCCTACATGCGCCCACTAGCCGTGGAGGGGCATCCGTGGAGGAGACCGACCAAAGCCACTTGGCCACTGCGGGCTCTACTTCATCGCATTCCTTGCAAAAGTATTACATCACGGGGGAGGCAGAGGGTTTCCCTGCCACAGGCCGGCCAGAATCGGGCGCCATGGGCTCTAGCCTGGACGACGAGCACATCCTGAGCGCCCTGCTGCAGAGCGACGACGAACTGGTGGGCGAGGACAGCGACAGCGAGGTCAGCGACCACGTGTCCGAGGACGACGTGCAGTCCGACACCGAGGAAGCCTTCATCGACGAGGTGCACGAAGTGCAGCCTACCAGCAGCGGCTCCGAGATCCTGGACGAGCAGAACGTGATCGAGCAGCCTGGCAGCTCCCTGGCCAGCAACAGAATCCTGACCCTGCCCCAGAGAACCATCAGAGGCAAGAACAAGCACTGCTGGTCCACCTCCAAGAGCACCAGGCGGAGCAGAGTGTCCGCCCTGAACATCGTGCGGAGCCAGAGGGGCCCCACCAGAATGTGCAGAAACATCTACGACCCCCTGCTGTGCTTCAAGCTGTTCTTCACCGACGAGATCATCAGCGAGATCGTGAAGTGGACCAACGCCGAGATCAGCCTGAAGAGGCGGGAGAGCATGACCAGCGCCACCTTCAGAGACACCAACGAGGACGAGATCTACGCCTTCTTCGGCATCCTGGTGATGACCGCCGTGAGAAAGGACAACCACATGAGCACCGACGACCTGTTCGACAGATCCCTGAGCATGGTGTACGTGTCCGTGATGAGCAGAGACAGATTCGACTTCCTGATCAGATGCCTGAGAATGGACGACAAGAGCATCAGACCCACCCTGCGGGAGAACGACGTGTTCACCCCCGTGCGGAAGATCTGGGACCTGTTCATCCACCAGTGCATCCAGAACTACACCCCTGGCGCCCACCTGACCATCGATGAGCAGCTGCTGGGCTTCAGAGGCAGATGCCCCTTCAGAGTGTACATCCCCAACAAGCCCAGCAAGTACGGCATCAAGATCCTGATGATGTGCGACAGCGGCACCAAGTACATGATCAACGGCATGCCCTACCTGGGCAGAGCCACCCAGACAAACGGCGTGCCCCTGGGCGAGTACTACGTGAAAGAACTGAGCAAGCCTGTGCATGGCAGCTGCAGGAACATCACCTGCGACAACTGGTTCACCAGCATCCCCCTGGCCAAGAACCTGCTGCAGGAACCCTACAAGCTGACCATCGTGGGCACCGTGCGGAGCAACAAGCGGGAGATCCCAGAGGTGCTGAAGAACAGCAGATCCAGACCTGTGGGAACAAGCATGTTCTGCTTCGACGGCCCCCTGACCCTGGTGTCCTACAAGCCCAAGCCCGCCAAGATGGTGTACCTGCTGTCCAGCTGCGACGAGGACGCCAGCATCAACGAGAGCACCGGCAAGCCCCAGATGGTGATGTACTACAACCAGACCAAGGGCGGCGTGGACACCCTGGACCAGATGTGCAGCGTGATGACCTGCAGCAGAAAGACCAACAGATGGCCCATGGCCCTGCTGTACGGCATGATCAATATCGCCTGCATCAACAGCTTCATCATCTACAGCCACAACGTGTCCAGCAAGGGCGAGAAGGTGCAGAGCCGGAAGAAATTCATGCGGAACCTGTACATGAGCCTGACCTCCAGCTTCATGAGAAAGAGACTGGAAGCCCCCACCCTGAAGAGATACCTGCGGGACAACATCAGCAACATCCTGCCCAAGGAAGTGCCAGGAACAAGCGACGACAGCACCGAGGAACCCGTGATGAAGAAGAGGACCTACTGCACCTACTGTCCCAGCAAGATCAGAAGAAAGGCCAACGCCAGCTGCAAGAAATGCAAAAAAGTGATCTGCCGGGAGCACAACATCGACATGTGCCAGAGCTGTTTCTCTAGGGCGGACCCAAAAAAGAAAAGGAAGGTGGATATCGCTGGAGACATGAGAGCTGCCAACCTTTGGCCAAGCCCGCTCATGATCAAACGCTCTAAGAAGAACAGCCTGGCCTTGTCCCTGACGGCCGACCAGATGGTCAGTGCCTTGTTGGATGCTGAGCCCCCCATACTCTATTCCGAGTATGATCCTACCAGACCCTTCAGTGAAGCTTCGATGATGGGCTTACTTACCAACCTGGCAGACAGGGAGCTGGTTCACATGATCAACTGGGCGAAGAGGGTGCCAGGCTTTGTGGATTTGACCCTCCATGATCAGGTCCACCTTCTAGAATGTGCCTGGCTAGAGATCCTGATGATTGGTCTCGTCTGGCGCTCCATGGAGCACCCAGTGAAGCTACTGTTTGCTCCTAACTTGCTCTTGGACAGGAACCAGGGAAAATGTGTAGAGGGCATGGTGGAGATCTTCGACATGCTGCTGGCTACATCATCTCGGTTCCGCATGATGAATCTGCAGGGAGAGGAGTTTGTGTGCCTCAAATCTATTATTTTGCTTAATTCTGGAGTGTACACATTTCTGTCCAGCACCCTGAAGTCTCTGGAAGAGAAGGACCATATCCACCGAGTCCTGGACAAGATCACAGACACTTTGATCCACCTGATGGCCAAGGCAGGCCTGACCCTGCAGCAGCAGCACCAGCGGCTGGCCCAGCTCCTCCTCATCCTCTCCCACATCAGGCACATGAGTAACAAAGGCATGGAGCATCTGTACAGCATGAAGTGCAAGAACGTGGTGCCCCTCTATGACCTGCTGCTGGAGGCGGCGGACGCCCACCGCCTACATGCGCCCACTAGCCGTGGAGGGGCATCCGTGGAGGAGACCGACCAAAGCCACTTGGCCACTGCGGGCTCTACTTCATCGCATTCCTTGCAAAAGTATTACATCACGGGGGAGGCAGAGGGTTTCCCTGCCACAGCTCCTGCAGGTGGAGGCTCTGCTGGAGACATGAGAGCTGCCAACCTTTGGCCAAGCCCGCTCATGATCAAACGCTCTAAGAAGAACAGCCTGGCCTTGTCCCTGACGGCCGACCAGATGGTCAGTGCCTTGTTGGATGCTGAGCCCCCCATACTCTATTCCGAGTATGATCCTACCAGACCCTTCAGTGAAGCTTCGATGATGGGCTTACTTACCAACCTGGCAGACAGGGAGCTGGTTCACATGATCAACTGGGCGAAGAGGGTGCCAGGCTTTGTGGATTTGACCCTCCATGATCAGGTCCACCTTCTAGAATGTGCCTGGCTAGAGATCCTGATGATTGGTCTCGTCTGGCGCTCCATGGAGCACCCAGTGAAGCTACTGTTTGCTCCTAACTTGCTCTTGGACAGGAACCAGGGAAAATGTGTAGAGGGCATGGTGGAGATCTTCGACATGCTGCTGGCTACATCATCTCGGTTCCGCATGATGAATCTGCAGGGAGAGGAGTTTGTGTGCCTCAAATCTATTATTTTGCTTAATTCTGGAGTGTACACATTTCTGTCCAGCACCCTGAAGTCTCTGGAAGAGAAGGACCATATCCACCGAGTCCTGGACAAGATCACAGACACTTTGATCCACCTGATGGCCAAGGCAGGCCTGACCCTGCAGCAGCAGCACCAGCGGCTGGCCCAGCTCCTCCTCATCCTCTCCCACATCAGGCACATGAGTAACAAAGGCATGGAGCATCTGTACAGCATGAAGTGCAAGAACGTGGTGCCCCTCTATGACCTGCTGCTGGAGGCGGCGGACGCCCACCGCCTACATGCGCCCACTAGCCGTGGAGGGGCATCCGTGGAGGAGACCGACCAAAGCCACTTGGCCACTGCGGGCTCTACTTCATCGCATTCCTTGCAAAAGTATTACATCACGGGGGAGGCAGAGGGTTTCCCTGCCACA
